# Fine-tuned protein language model identifies antigen-specific B cell receptors from immune repertoires

**DOI:** 10.1101/2025.10.30.685465

**Authors:** Karen Paco, Mariana Paco Mendivil, Zihao Zhang, Sanaz Zebardast, Christian Davila, Ryan M. Mooney, Peace Olatoyinbo, Tristan Yang, Sebastian Bassi, Virginia Gonzalez, Eva Chen, Faisal Bin Ashraf, Isabel Condori Roman, Jonathan R. Felix, Rashid M. Alam, Jordan A. Lay, Malkiat S. Johal, Karine G. Le Roch, Ilya Tolstorukov, Jeniffer B. Hernandez, Fernando L. Barroso da Silva, Stefano Lonardi, Matthew H. Sazinsky, Animesh Ray

**Affiliations:** Henry E. Riggs School of Applied Life Sciences, Keck Graduate Institute, Claremont CA, USA; Pomona College, Claremont, CA, USA; Department of Computer Science and Engineering, University of California, Riverside, CA, USA; Department of Molecular, Cell and Systems Biology (MCSB), University of California, Riverside, CA, USA; Universidade de São Paulo, Brazil; Department of Chemical and Biomolecular Engineering, NC State University, Raleigh, NC, USA; Toyoko Labs, Toyoko LLC, Emeryville, CA, USA; Division of Biology and Biological Engineering, California Institute of Technology, Pasadena, CA, USA

**Keywords:** Antibody Discovery, Computational Antibody Discovery, Antigen Specificity, Immune Repertoire, Protein Language Model, Antibody Diversity, Antibody Heterogeneity

## Abstract

Scalable identification of antigen-specific antibodies from whole immune repertoire V(D)J sequences is a central challenge in biomedical engineering. We show that protein language models (PLMs) fine-tuned on antibody heavy-chain sequences can directly predict antigen specificity from unselected immune repertoires. We assessed our model, Antigen Specificity Predictor (ASPred), against SARS-CoV-2, influenza, and HIV-AIDS antigens, observing comparable predictive performance. In the whole immune repertoire V(D)J sequences of mice immunized with the SARS-CoV-2 spike protein’s receptor-binding domain (RBD), ASPred identified antibody sequences specific to RBD. Several candidate sequences were validated, including one as a heavy chain-only nanobody with 20.7 nM dissociation constant. Molecular dynamics simulations supported the predicted interactions at coarse-grained and atomic levels. Benchmarking against Barcode-Enabled Antigen Mapping (BEAM) of B cell receptor sequence data had highly significant overlaps with ASPred predictions, suggesting scalability. The predicted SARS-CoV-2 binders differed substantially from training sequences, demonstrating generalization beyond sequence memorization. Together, we establish that heavy chain antibody sequences encode sufficient information for PLMs to infer specificity, offering a scalable framework for antibody discovery with broad applications.

## 1 Introduction

Adaptive immunity depends on mutation and recombination in the heavy (H) and light (L) chain genes in T and B cells [1–3], generating a large diversity of potential antigen-binding receptors [4, 5]. B lymphocytes (B cells) that express antigen-specific B cell receptors (BCRs) are selected through the interaction of the latter with surface-displayed antigen fragments; B cells then mature through further somatic mutagenesis and signaling events [6, 7]. Upon activation, selected B cells clonally expand and may undergo immunoglobulin class switching, a recombination process that replaces the heavy chain constant region with a different isotype (e.g., IgG, IgA, or IgE) to engage different aspects of the immune system function while maintaining the same variable regions and thus the same antigen specificity [8].

Our ability to identify antigen-specific antibodies from the whole immune repertoire is crucial for understanding immune responses and for accelerating counter-measure development against emerging pathogens. Experimental approaches focus on isolating/identifying individual antibodies, such as monoclonal antibodies (mAb) through hybridoma technology, surface displays, or by single BCR or T cell receptor sequencing [9]. Rapid sequencing of genes that encode circulating antibodies after SARS-CoV-2 virus infection led to an unprecedented speed in the production of mAbs against the COVID-19 pandemic [10–12].

The human antibody repertoire theoretically exceeds 10^16^ unique sequences [5, 13, 14], though far fewer are belived to be expressed. This diversity arises through V(D)J recombination, involving DNA rearrangement and mutation in the antibody gene segments, especially within the complementarity-determining regions (CDRs) [8, 15]. These variants undergo selection for improved antigen binding [16], however, the structural basis of B-cell clone selection and maturation is not fully understood [17, 18]. Only a small fraction (0.01–0.1%) of B cells in an immunized individual target a specific antigen, and mapping epitopes to antibody binding sites remains challenging.

High throughput experimental methods such as antigen-specific B cell sorting and BCR sequencing can identify reactive sequences [9, 19], yet these methods are technically demanding, expensive, and provide limited insight into the structural basis of antibody specificity [7, 20]. These constraints motivate computational strategies that can work directly from unselected immune repertoire sequences and ask whether one can predict specificity by learning short- and long-range protein sequence relationships common to a class of antibodies specific against an antigen. Models like Antiformer and AntiBERTa are examples of notable progress [21, 22], but neither platform classifies antigen-specific antibodies from the whole immune repertoire.

The emergence of protein language models (PLMs) trained on millions of natural protein sequences to learn both short- and long-range dependencies opened a new frontier [23–25]. By fine-tuning PLMs by task-specific supervision, it should be possible to transfer the general biochemical information about specific antigen-antibody binding pairs and their associated features to specialized tasks, including repertoire-scale classification and identification of antigen-specific BCRs from unpaired sequences— precisely where labeled datasets are scarce and high-throughput scalability is important. One such study trained a memory B-cell language model (mBLM) for chosen influenza antibodies for their respective binding epitope discovery [26]. To computationally predict binding affinities of antibodies to their preselected epitopes and to further optimize antibody sequences, a transformer model, AlphaBind, used PLM embeddings [27]. A sequence-based PLM, MAGE, fine-tuned to generate paired human heavy and light chain variable regions conditioned on an antigen sequence, successfully optimized experimentally verifiable, novel antibodies [28]. Similarly, we reported Ab-Affinity, an LLM integrated with combinatorial optimization algorithms (genetic algorithm and simulated annealing) to iteratively refine mutated antibody sequences toward higher predicted binding affinity [29]. Recently, a novel antibody language model pretrained on antigen specificity tasks significantly outperformed models using frozen embeddings [30], though novel predicted antibodies remained to be validated. These successes suggest that signals exist in antibody structures and their cognate antigens recognizable by deep neural networks.

While PLM based models have made significant advances in specificity determination, a critical gap remains in scaling and generalizability in performing high-throughput, repertoire-scale identification of antigen-specific antibodies directly from unpaired heavy-chain sequence data alone. Here we report a novel tool, Antigen Specificity Predictor or ASPred (Fig. 1), a specific-purpose PLM framework trained and fine-tuned on antibody heavy chain sequences, which can identify with high confidence antibodies specific to an antigen at the whole immune repertoire scale, validated on both human and mouse experimental data, and generalized to three different antigens: SARS-CoV-2 spike protein receptor binding domain (RBD), influenza hemagglutinin antigen (HA), and human immunodeficiency virus (HIV) antigen gp120.

**Fig. 1:**
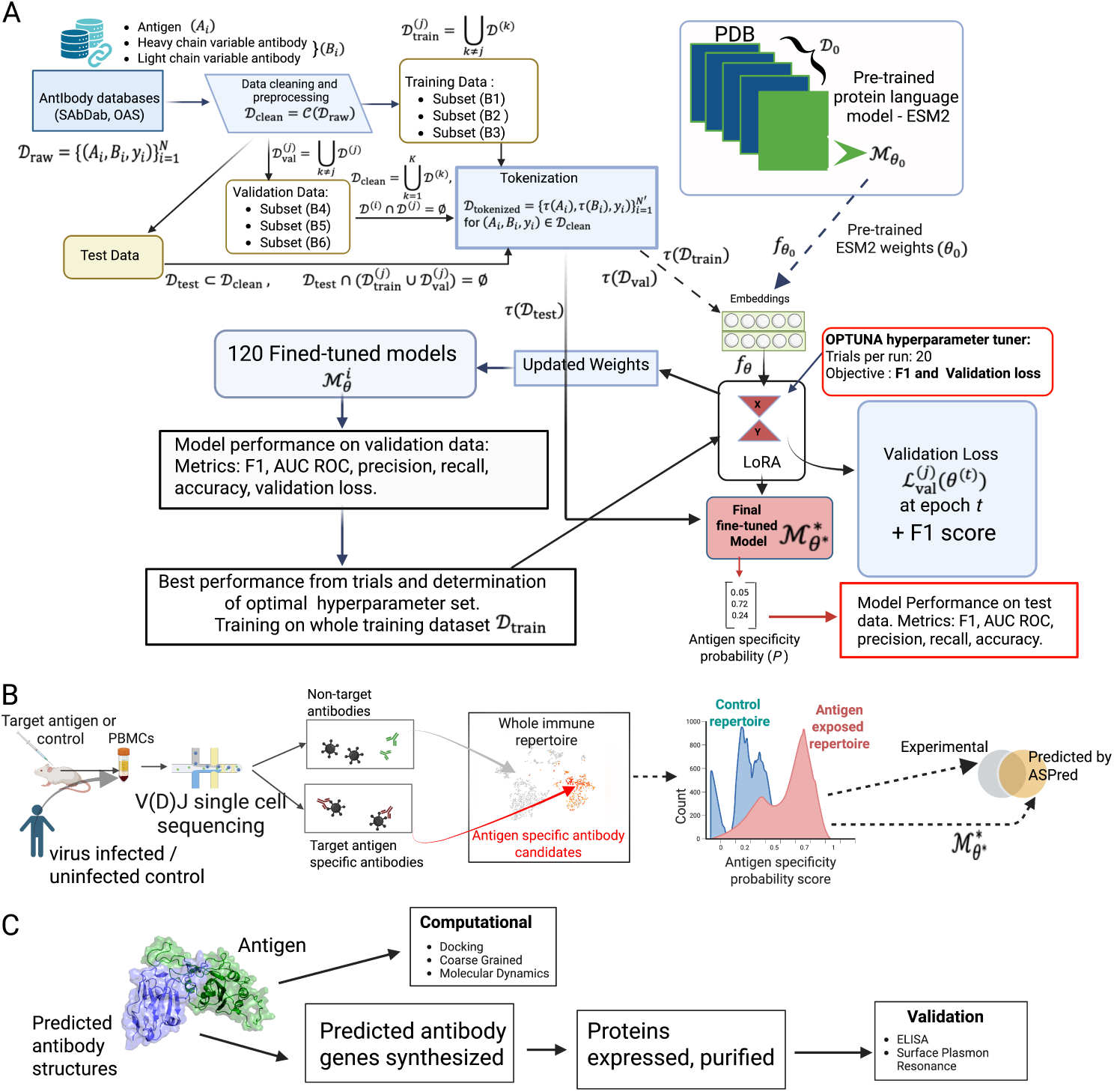
The strategy for constructing and validating ASPred. **(a)**, Schematic of fine-tuning ESM-2 to develop ASPred. Heavy chain antibody sequences (with data partitions provided in the diagram) were embedded using the ESM-2 PLM, then fine-tuned via parameter-efficient LoRA adaptation. A classification head outputs the antigen specificity probability *P_s_* for each antibody sequence. **(b)**, Validation of ASPred predictions applied to representative immune repertoire data and benchmarking against experimental data. RBD-immunized mouse whole immune repertoire was obtained by V(D)J sequencing of barcoded single cells without affinity tagging with the antigen. Antigen specificity scores by ASPred were obtained, and the resulting distributions enabled prioritization of high-confidence binders over non-binders. Human V(D)J sequence data from SARS-CoV-2 infected patients with RBD specificity scores from Barcode Enabled Antigen Mapping (BEAM) were used for ASPred inference testing. The same V(D)J sequence datasets were scored and stratified by ASPred according to predicted specificity. The overlap between the experimentally observed and predicted antibodies and their frequency distributions were compared. Publicly available human SARS-CoV-2 infected whole immune repertoires (and healthy controls) sequence data were used for teh inf_4_erence of RBD-specificity with ASPred. **c**, Targeted validation of ASPred-predicted antigen specificity of antibodies in the whole immune repertoire (above) by gene synthesis, cloning, protein expression, purification, binding assays against the antigen, and molecular dynamics simulation.

## 2 Results

Using publicly available paired antigen-antibody sequence data (Fig. 1a), we fine-tuned the general embeddings of a PLM to high predictive accuracy (ASPred), with strong performances on three different antigen classes, explored its predictions on previously unseen data, used it to make predictions to classify novel antigen-specific antibodies in mice whole-immune repertoire by single cell V(D)J sequencing and high throughput experimentally obtained antigen-specific human antibody sequences (Fig. 1b), validated the candidates by expressing in *E. coli* as nanobodies (Fig. 1c), and examined its performance on unseen human whole immune repertoires.

### 2.1 Heavy-chain sequences enable accurate antigen-specificity classification

We began with the pretrained general protein language model ESM-2 backbone to predict antigen specificity from heavy-chain sequences. We fine-tuned all ESM-2 parameters end-to-end using a randomly initialized classification head (see Methods). Our first model (Model 1), a fully fine-tuned ESM-2, reached *>* 80% accuracy on a held-out SARS-CoV-2 RBD heavy-chain test set (Supplementary Fig. S1). By contrast, an antibody-specific model (AbLang embeddings[31] + logistic regression), achieved 55.2% (±0.0091) on this benchmark (Extended Data Fig. 1). We subsequently performed parameter-efficient fine-tuning (LoRA) of ESM-2 backbones with 8M and 650M parameters (Fig. 1a) (see descriptions of training datasets in Extended Data; their partitions are diagrammed in (Fig. 1a) and in Methods). Trained on antibodies (heavy, light, and heavy + light chains) against three different antigens, namely, SARS-CoV-2 RBD, influenza HA, and HIV glycoprotein gp120, respectively, we generated two models for each antigen–ASPred F1 (F1-optimized) and ASPred VL (validation loss optimized). These six ASPred models outperformed ESM-2 (Fig. 2a–c).

**Fig. 2:**
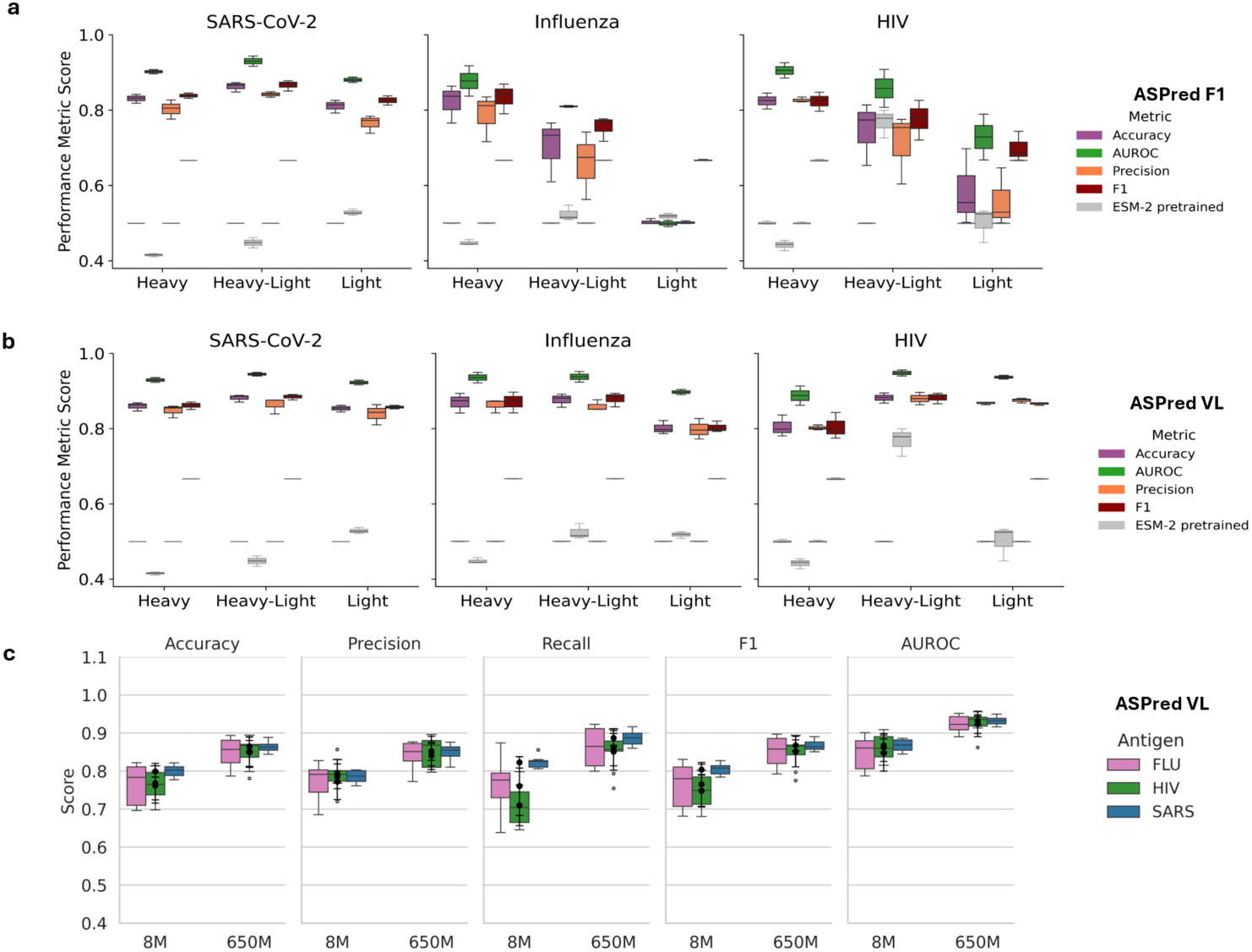
Performance evaluation of ASPred models across three antigens. (**a**,**b**) Cross-validated performance (accuracy, AUROC, precision, F1) for SARS-CoV-2 RBD, influenza HA, and HIV gp120 using heavy, light, and paired heavy–light-chain inputs; (**a**) F1-optimized models, (**b**) Validation loss-optimized models, (**c**) Performance fine-tuning of 8M and 650M parameter ESM-2 models with heavy chains (metrics: accuracy, precision, recall, F1, and AUROC. The model configurations are summarized in the Supplementary Table 1. Dataset sources and selected hyperparameters are detailed in Extended Data, Methods, and Supplementary Results.

To examine the learned representations, we projected the heavy chain sequence embeddings into two dimensions using t-SNE (Figure 3a,b) and two other dimensionality reduction methods (see Extended Data Table 1). Embeddings from the fine-tuned models displayed statistically significant improvement in separation between antigen binders and non-binders across all three antigen types relative to the pre-trained baseline as determined by silhouette score [32] and the Davies–Bouldin index [33](Extended Data Table 1).

**Fig. 3:**
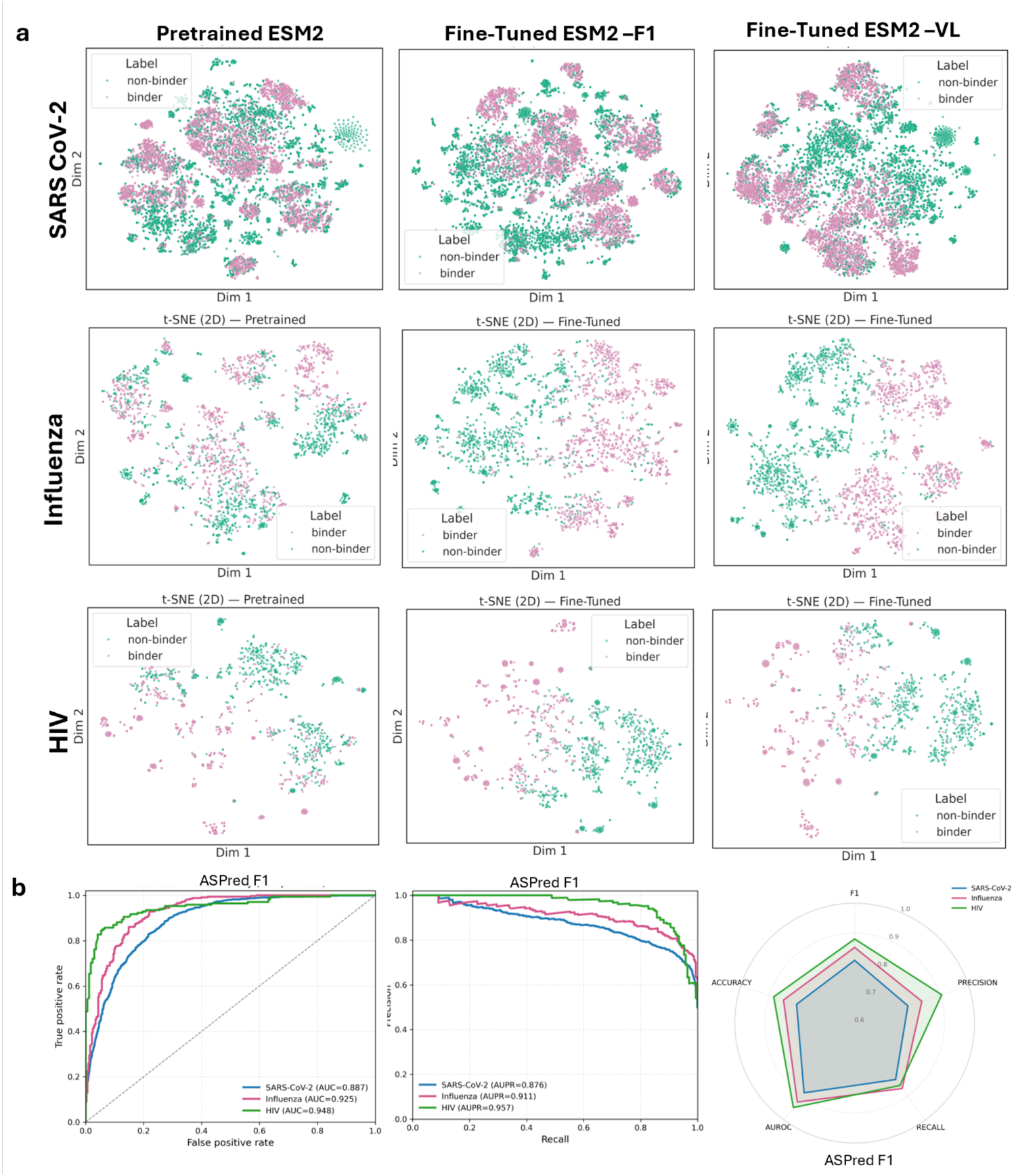
Differences in sequence embeddings upon fine-tuning of ESM-2 to ASPred F1 and VL models across three antigens. (**a**) Two-dimensional t-SNE projections of sequence embeddings before fine-tuning (pretrained) and after fine-tuning (ASPred-F1, ASPred-VL). Purple, BCR sequences predicted to be antigen specific; green, predicted to be not specific to the given antigen. (**b**) Receiver Operating Characteristic (ROC) curve,) Precision Recall (PR) and Radar plot summarizing performance metrics for the final model ASPred-F1, evaluated on an independent held-out test set (see Supplementary Fig. S2)for ASPred VL).

### 2.2 ASPred classifies unseen human RBD-specific BCRs

We assessed the predictive performance of ASPred against an established high-throughput experimental method for directly identifying antigen-specific BCRs, namely Barcode Enabled Antibody Mapping (BEAM) technology [34]).

The BEAM score (0-100; rank-based) provides a quantitative measure of antigen specificity. ASPred, by contrast, outputs a per-sequence probability (*p ∈* [0, 1]) of antigen binding. Unless noted otherwise, we considered sequences with *p ≥* 0.5 to be antigen-specific (AS) (see Supplementary Figure). Although neither metric directly measures binding affinity, both serve as useful proxies for identifying putative AS BCRs. On a previously unseen BEAM dataset of RBD-specific BCR sequences (10x Genomics), ASPred predictions showed highly significant overlap (Figure 4a); total BCR sequences, *n* = 2,464; *p* = 8.5 *×* 10*^−^*^66^; odds ratio (OR) = 15.9 using ASPred F1; and *p* = 7.49 *×* 10*^−^*^68^; OR = 14.8 using ASPred VL). However, their concordance was moderate (Jaccard Index 0.33), which is not unexpected given that the two methods are very different: one is based on a high-throughput microfluidic *in vitro* ligand-binding assay (BEAM) and the other is an entirely computational prediction system (ASPred). Nonetheless, the overlap was significant.

**Fig. 4:**
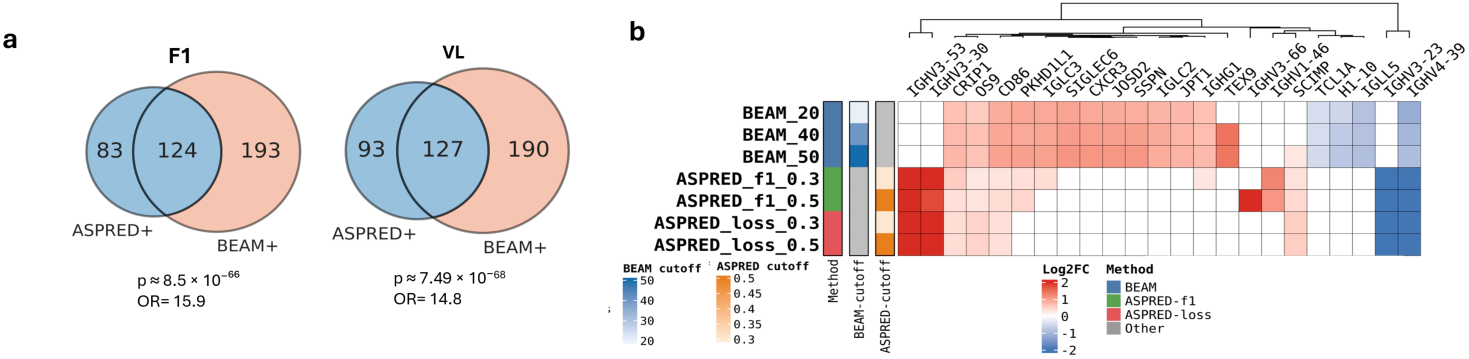
Benchmarking against BEAM. (**a**) Overlap between sequences predicted by ASPred (ASPred^+^) and experimentally discovered by BEAM technology (BEAM^+^) at fixed thresholds for ASPred-F1 and ASPred-VL (Total candidate BCR sequences selected randomly: *n* = 2,464). (**b**) Concordance between upregulated gene signatures in cells expressing the corresponding BCR V(D)J sequences across BEAM and ASPred thresholds (hierarchically clustered heat map of enrichment scores). The data are from 10x Genomics scRNAseq and V(D)J sequencing, from sorted B cells and challenged with RBD. BEAM scores of 20, 40 and 50 were used as cutoffs for admitting cells as antigen-specific (BEAM+); similarly, the same BCR sequences from V(D)J reads had ASPred scores with thresholds of 0.3 and 0.5. The mRNA expression level was measured as log-normalized UMI counts.

We then asked how different are the gene expression patterns of AS B cells relative to those of B cells that are not antigen-specific (ANS) as defined by either ASPred prediction or by BEAM experiments. We linked V(D)J heavy-chain sequences to their corresponding single cell RNA deep sequencing data (scRNA-seq). We were thus able to perform differential mRNA expression analysis between AS and ANS B cells for each of the two platforms (see Methods).

The heatmap in (Figure 4b) shows differential expression (DE) log_2_ FC (log fold change) across ASPred/BEAM thresholds. This analysis suggests that ASPred-predicted AS cells exhibit transcriptomic signatures aligned with prior observations even when they do not perfectly overlap with BEAM results. In antigen-specific cells predicted by ASPred, IGHV3-53 transcript levels are up-regulated. This is consistent with previous reports of upregulation of this transcript in B cells in blood containing SARS-CoV-2–neutralizing antibodies [35–37] (Figure 4b). Furthermore, the gene for the adapter protein SCIMP, a transmembrane scaffold for TLR4 that drives IL-6 production in macrophages and is expressed in B cells and other professional antigen-presenting cells at the immunological synapse via tetraspanin-enriched microdomains, has been implicated in COVID-19 lung hyper-inflammation and host-defense programs [38]. We also observed, in congruence with the previous observations in longitudinal Omicron BA.2 breakthrough-infection cohorts, a down-regulation of CRIP1 in naïve B cells [39]. The divergence between BEAM-defined and ASPred-defined differential gene expression signatures likely represent overlapping yet somewhat distinct cell populations, reflecting their different methodological origins: experimental rank-based *in vitro* binding calls in BEAM technology versus *in silico* sequence-derived specificity probabilities of ASPred.

### 2.3 ASPred identified at high frequency RBD-specific antibodies in murine repertoire

To evaluate ASPred’s performance, we applied it to *de novo* BCR V(D)J heavy-chain repertoire sequences without affinity selection from mice immunized with the SARS-CoV-2 RBD (2019-nCoV) antigen (Figure 5a; see Methods and Extended Data). From 312 unique heavy chain sequences identified, ASPred Model 1 classified 30 AS BCR candidates (Figure S7).

**Fig. 5:**
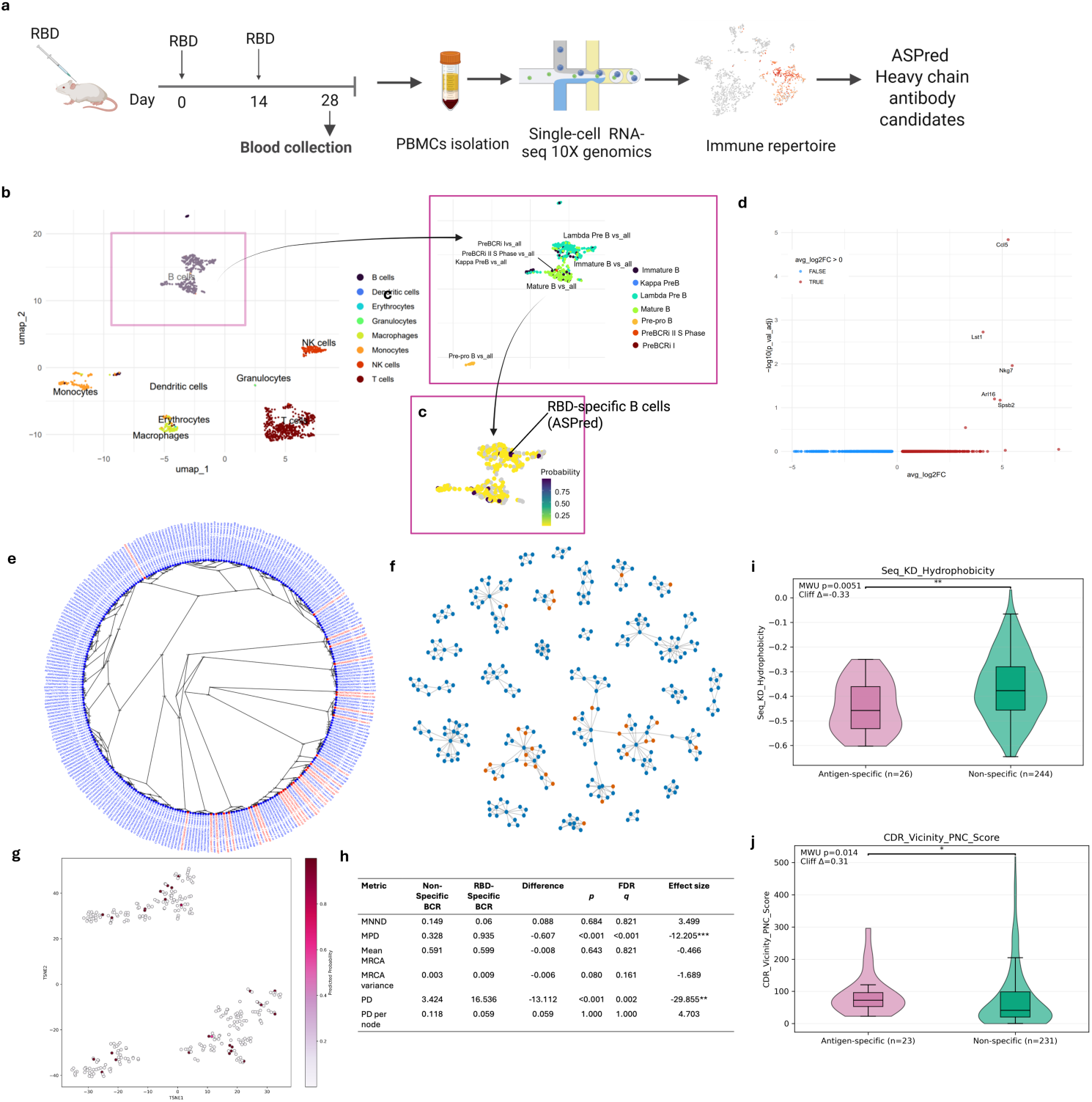
Integrated single-cell transcriptome and repertoire analysis of antigen-specific B cells. (**a**) Experimental workflow: Mice were immunized, and single peripheral blood mononuclear cells (PBMC) were isolated by 10x Genomics Chromium chip, barcoded and scRNAseq/V(D)J sequences were obtained, filtered and analyzed as described in Methods. The V(D)J heavy sequences were analyzed by ASPred to provide a specificity score. (**b**) UMAP of RBD immunized whole immune repertoire from Peripheral Blood Mononuclear Cells (PBMCs) (**c**) Overlay of RBD-specific cells and their ASPred specificity scores is shown by the color spectrum. (**d**) Differential gene expression analysis of RBD-specific and non-specific B cells from the whole immune repertoire. (**e,f**) Maximum likelihood tree alignment of the VHH sequences of the whole immune repertoire B cells (100 bootstraps). V(D)J sequences were trimmed (see Methods) (e) and K-nearest neighbor (k=3) network (f) of the AS and ANS BCR sequences. Edges represent the nearest neighbors. Red, AS BCRs; blue, ANS BCRs (see Methods). (**g**) Low-dimensional sequence-space embedding showing AS BCRs (magenta) and ANS BCRs (gray) dispersed across clusters. (**h**) Measures of phylogenetic diversity metrics of the B cell receptor repertoire. We measured phylogenetic sequence diversity by mean near^1^e^0^st-neighbor distances (MNND), mean and variance of the node-to-most-recent common ancestor (MRCA) distances, patriarchal distances (PD), and PD per node (see Methods). (**i**) Violin + box plots of sequence hydrophobicity (Kyte–Doolittle GRAVY); distributional shift between AS BCRs and ANS BCRs (two-sided Mann–Whitney *U*; Extended Data). (**j**) Violin + box plots of CDR-vicinity Patches of negative charge (PNC); AS BCRs differ from ANS BCRs (two-sided Mann–Whitney *U*; exact *P* in Extended Data).

### 2.4 Predicted antigen-specific B cells resembled but are not identical to known SARS-CoV-2 stimulated B cells

We asked which B-cell subsets carry ASPred-predicted RBD-specific BCRs. Using paired V(D)J sequences and scRNA-seq of the murine PBMC repertoire. We annotated immune lineages by B-cell subtypes [40] (Figure 5b). Most B cells displaying V(D)J sequences classified as RBD-specific by ASPred were annotated as mature B cells (Figure 5c), with a minor fraction labeled as lambda-expressing pre–B-like cell (Extended Data). Because pre-B cells are uncommon in PBMCs, we interpret this latter annotation cautiously as likely to be a reference-mapping artifact (Extended Data).

Next, we tested whether mature B cells that displayed ASPred-predicted RBD-specific BCRs exhibit distinct transcriptional states relative to other immune cell types. Differential gene expression analysis between AS B cells to ANS B cells in the murine repertoire identified five genes: *CCL5*, *NKG7*, *LST1*, *ARL16* and *SPSB2* (Figure 5d). Upregulation of *CCL5*, encoding a chemokine that promotes T-cell recruitment and activation, is consistent with enhanced T and B cell collaboration during B cell activation. Upregulation of *NKG7* and *LST1* suggest recent immune activation. Full statistics and gene lists are provided in Supplementary Tables. Together, these results indicate that ASPred enriches for B cells exhibiting transcriptional hallmarks of antigen experienced B cells and T-helper cells.

### 2.5 ASPred-predicted mouse VHH antibodies against RBD exhibit high sequence diversity

We examined whether ASPred predicted AS antibodies might be biased towards one sequence class, which might be expected if the ASPred LLM had memorized certain sequences. By contrast, multiple sequence alignment (MSA) of their heavy chains highlighted diverse CDRH3 motifs (Fig. 5e). K-nearest neighbor (kNN) graph (Fig. 5f) of the combined BCR sequences of the whole murine repertoire confirmed this conclusion, but also demonstrated that while the predicted RBD-specific BCRs were more clonal than others, they also represented diverse clonotypes. The predicted BCRs against RBD were broadly dispersed across multiple clusters (Fig.3g), indicating substantial sequence diversity (see t-SNE parameter settings in Methods) among the ASPred-predicted AS antibodies. Analysis of phylogenetic diversity metrics (Fig.3h) revealed a clear contrast between AS and ANS BCR sequences: RBD-specific sequences exhibited substantially lower mean pairwise distance (MPD) and total phylogenetic diversity (PD) compared with ANS sequences (both q < 0.01), indicating that antigen-specific clones occupy a restricted region of the sequence space and share recent ancestry. The relatively compact topology of the KNN graph (Fig. 5f) and the relatively dispersed AS sequence clusters are consistent with clonal expansion and convergent affinity maturation following antigen exposure. In contrast, ANS BCR sequences displayed a much broader spread across the tree, were less clonal and were more dispersed in the tSNE plots, reflecting a heterogeneous, unselected background repertoire. Other metrics, including mean nearest-neighbor distance, mean and variance of node-to-most recent common ancestor distances, and PD normalized per node, did not differ significantly between the two groups, suggesting that local branching structures within each subset are similar even though ASPred-predicted RBD-specific lineages clustered tightly within the larger repertoire. Overall, these patterns demonstrate that the predicted RBD-specific sequences are phylogenetically constrained and less diverse, consistent with selective focusing of the B-cell response toward the immunizing antigen. Finally, we asked whether ASPred-predicted RBD-specific BCR sequences from mouse whole immune repertoires exhibit distinct biophysical properties relative to non-specific antibodies by quantifying three CDR-proximal surface metrics: (i) CDR-vicinity patch of negative charges; (ii) CDR-vicinity patch of positive charges; and (iii) CDR-vicinity hydrophobic patch score [41, 42]. We also evaluated several sequence-level properties, namely, Kyte-Doolittle GRAVY, isoelectric point, and net charge at pH 7 for these two sets of antibodies. Relative to ASPred-predicted RBD-non-specific BCRs, RBD-specific BCRs showed higher hydrophobicity (Kyte–Doolittle GRAVY; *P* = 0.0051, *δ* = 0.33; Fig. 5i), and higher PNC (two-sided Mann–Whitney *U*, *P* = 0.014, Cliff’s *δ* = 0.31; Fig. 5j). Other parameters did not exhibit significant difference (upon correction for multiple testing) (see Extended Data). The higher than average surface negatively charged patches and higher hydrophobicity in the predicted RBD-specific antibodies are consistent with the presence of a high net negatively charged surface area (contributed by E484, D405, D420, and D467) localized as a patch, and a significant hydrophobicity in the RBD domain (contributed by F486, Y489, L455, and Y505) over the surface of interaction with ACE2.

### 2.6 ASPred-predicted nanobodies bind to RBD *in vitro*

We synthesized and recombinantly expressed in *E. coli* as nanobodies ten randomly chosen murine RBD-specific variable heavy domain of heavy chain (VHH) antibody candidates exhibiting ASPred probability score *p ≥* 0.5 alongside a known RBD-specific VHH C5 nanobody (PDB: 7OAO) [43]. Periplasmic proteins were tested for VHH binding to RBD by Western blot (Figure 6a), by enzyme linked immunosorbent assays (ELISA) (Figure 6b). Five of the ten candidates exhibited robust binding relative to the negative control (the remaining 5 did not show consistent binding). Ab-157 and Ab-200 were further purified (Supplementary Fig. S3) and again were confirmed for binding by ELISA assays (Supplementary Figure S4). Localized surface plasmon resonance (LSPR) confirmed Ab-157 binding to immobilized RBD with high affinity, *K_D_* = 20.7 nM, *k*_off_ = 6.72 *×* 10*^−^*^5^ s*^−^*^1^, and *k*_on_ = 3.253 *×* 10^3^ M*^−^*^1^s*^−^*^1^ (Fig. 6c,d). A binding signal was observed for Ab-200 as well, but non-specific interactions on the reference channel precluded definitive LSPR interpretation.

**Fig. 6:**
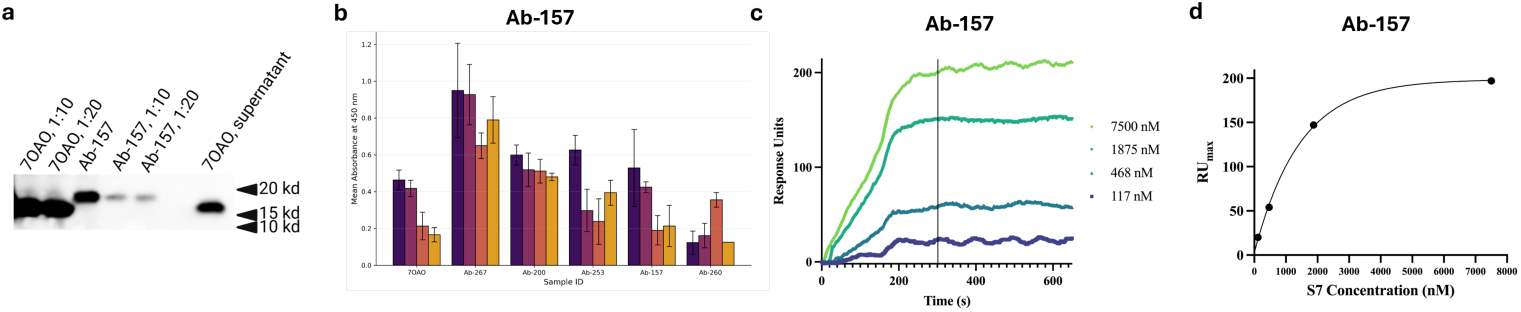
Experimental validation of ASPred-predicted antibodies. (**a**) Western blot confirming expression of selected VHHs. (**b**) ELISA binding to SARS-CoV-2 RBD for ten randomly chosen VHH candidates from ASPred outputs across serial dilutions (mean ± s.e.m.). (**c,d**) LSPR sensorgrams for purified VHHs; Ab-157 shows high-affinity binding (*K_D_* = 20.7 nM).

### 2.7 Dynamical simulations revealed interacting residues between RBD and its targeted VHH nanobodies predicted by ASPred

We implemented a multi-scale *in silico* validation pipeline integrating molecular docking, constant-pH coarse-grained (CG) Monte Carlo simulations, and atomistic constant-charge Molecular Dynamics (MD). VHH candidates were first folded and docked against the SARS-CoV-2 RBD (2019-nCoV) using IgFold [44] and ClusPro 2.0 [45] (Fig. 7). Ten predicted RBD-specific BCRs including Ab-157 and Ab-200 were selected. Multiple conformations were generated per antibody–antigen pair, and the top cluster structures determined by ClusPro were used to estimate binding energies using PRODIGY [46]. For comparison, we used the nanobody C5 (PDB ID 7OAO), a well-characterized nanobody binder of the RBD, as a positive control [47]. In congruence with the ELISA assay results, several candidate pairs demonstrated binding affinities comparable to or exceeding that of C5 to RBD as determined by PRODIGY(Supplementary Figure S7).

**Fig. 7:**
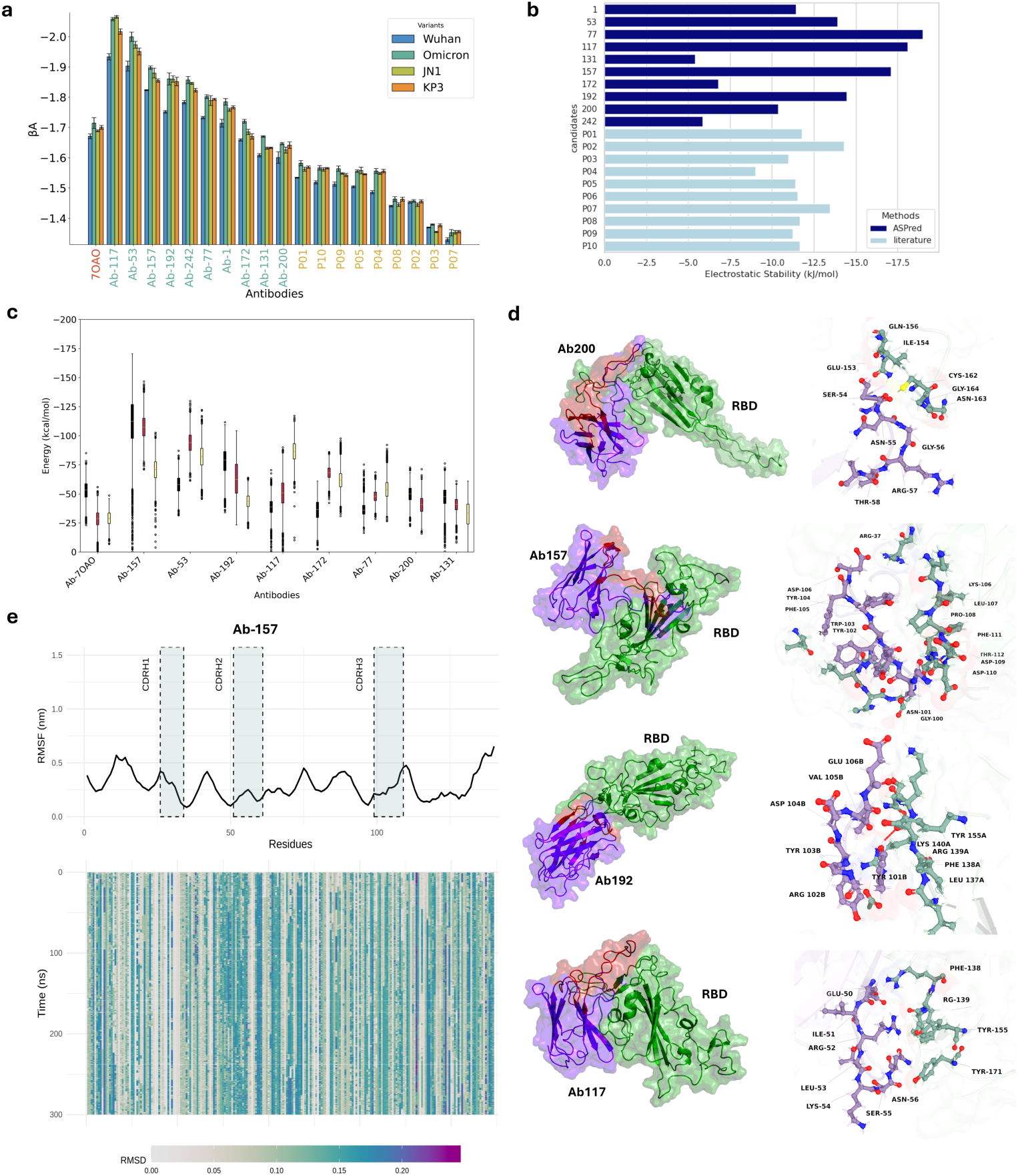
Multiscale *in silico* validation of ASPred-predicted VHH–RBD interactions. (**a**) Constant-pH coarse-grained simulations estimating affinity scores (*β_A_*) for four RBD variants (Wuhan, Omicron, JN.1, KP.3); bars show mean ± s.e.m. (**b**) Electrostatic stability of ASPred-predicted VHHs. (**c**) Binding free energies from explicit-solvent MD using MM-PBSA for predicted and positive control (Ab-70A0) for binding to the Wuhan RBD only; box plots summarize replicate trajectories per antibody. (**d**) Representative complexes for four antibodies (Ab192, Ab200, Ab117, Ab157). Left: RBD (green surface) with VHH (purple cartoon; CDRs in red). Right: interface zooms highlighting recurrent hydrogen bonds and salt bridges. (**e**) Stability analysis for Ab-157 across 300 ns MD. Top: per-residue RMSF with CDR windows indicated; bottom: residue-wise RMSD hea1t4map over time showing a stable paratope– epitope interface.

To evaluate interaction strength under physiologically relevant conditions and to include the solution pH effect in the calculation, we used the FORTE constant-pH CG simulation framework [48, 49] to compute free energies of binding at pH 7.4 and 1:1 salt concentration. All VHHs exhibited favorable binding profiles, consistent with values reported for potent monoclonal antibodies. As shown in Fig. 7a, ASPred candidates outperformed previously published antibody designs [50], which were produced via physics-based modeling approaches. Among them, Ab-117, Ab-53, and Ab-157 scored the highest in predicted binding affinities (Fig. 7a). Interestingly, candidates predicted by ASPred also exhibited stronger binding affinities for multiple SARS-CoV-2 variants, including Omicron BA.1 and more recently circulating strains such as JN.1 and KP.3. This might indicate that ASPred is capturing a broad reactivity across antigenic landscapes, both for the original antigenic protein and later emerging variants.

To further evaluate molecular stability, we applied the “Fast Proton Titration Scheme” (FPTS) [51] to calculate electrostatic stability based on the averaged net charge of titrable residues at physiological pH and salt conditions. ASPred-predicted antibodies displayed electrostatic stability comparable to or exceeding known anti-SARS-CoV-2 RBD antibodies ((Fig. 7b)), with Ab-117 again emerging as a top candidate due to its combined affinity and stability profile.

Finally, to probe fine-grained interactions at atomic resolution, we selected eight ASPred-predicted VHHs for explicit-solvent MD simulations in complex with the SARS-CoV-2 RBD (Fig. 7c). To quantify the strength of these interactions, binding free energies were computed using the Molecular Mechanics/Poisson-Boltzmann Surface Area (MM-PBSA) method. Across three independent replicates, ASPred-predicted VHHs exhibited similar or stronger affinities than the positive-control nanobody C5. Throughout 300 ns trajectories, VHH–RBD complexes remained conformationally stable with lower root mean squared deviation (RMSD) and residue flexibility (RMSF). Interface contact analyses showed persistent CDR-mediated interactions at RBD epitopes (Figure 7d; Extended Data). Among these Ab-157, the nanobody experimentally confirmed by LSPR to bind RBD, maintained a compact and persistent interface over the full 300 ns window (Fig. 7d). MM-PBSA per-residue decomposition highlighted strong contributions from CDRH3 residues, consistent with their frequent hydrogen bonding and electrostatic contacts at the paratope–epitope interface (Fig. 7e).

### 2.8 ASPred can identify RBD-specific BCRs from bulk sequencing data of whole immune repertoires

We collected heavy-chain V(D)J repertoire datasets from publicly available bulk PBMC sequences of healthy donors and SARS-CoV-2–infected individuals (dataset details in Extended Data). Within each repertoire, sequences were stratified into RBD antigen-specific BCRs (RBD AS-BCRs; AS^+^) and RBD non-specific BCRs (RBD ANS-BCRs; AS^-^) by applying a fixed ASPred probability threshold of *≥* 0.5. BCR diversity was quantified per sample and in pooled analyses using (i) Shannon entropy (*H*) [52, 53] and (ii) inverse Simpson (1/*D*) [54] computed on CDRH3-defined sequences. To control for depth differences, we used equal-size resampling (See methods).

Across repertoires, AS^-^ exhibited consistently higher diversity than AS^+^ by both *H* and 1/*D* (Figure 8a), suggesting that predicted antigen-specific sequences are concentrated in fewer clonotypes relative to the background. Concordantly, clonotype-size distributions showed a heavier tail in AS^-^, with a greater fraction of large clone sizes (e.g., *≥* 20 members), whereas AS^+^ was skewed toward smaller clone sizes (Fig. 8b). This pattern suggests that the ASPred^+^ calls capture focused sequence subsets within each repertoire rather than capturing broad and diffuse diversity. Since the information of diversity does not directly capture phylogenetic relationships, we also generated 100 partially synthetic datasets each comprising the same 127 antigen-specific AS^+^ and 127 randomly sampled AS^-^. We then generated 100 different phylogenic trees (100 bootstraps each). The median Fitch parsimony Z-scores showed a consistent deviation from the permutation null (Supplementary Figure S6a), indicating that the AS^+^ sequences were significantly more phylogenetically clustered than expected by chance. When all trees were pooled (Supplementary Figure S6b), the Z-score distribution was unimodal and sharply centered near −13, confirming robust AS^+^ clustering across replicates. The AS^-^-only clade size distribution (Supplementary Figure S6c) was right-skewed, with many small clades (3–6 tips) and progressively fewer large clusters, suggesting localized enrichment rather than global segregation. Correspondingly, the local AS^+^ enrichment metric (mean fraction of AS^+^ among k = 3 nearest neighbors; (Supplementary Figure S6d)) was tightly distributed around 0.9, indicating that most AS^+^ sequences have predominantly AS^+^ neighbors within the tree topology. Together, these results show that predicted antigen-specific antibody sequences form statistically significant, spatially localized clusters in phylogenetic space, consistent with convergent sequence features linked to antigen recognition.

**Fig. 8:**
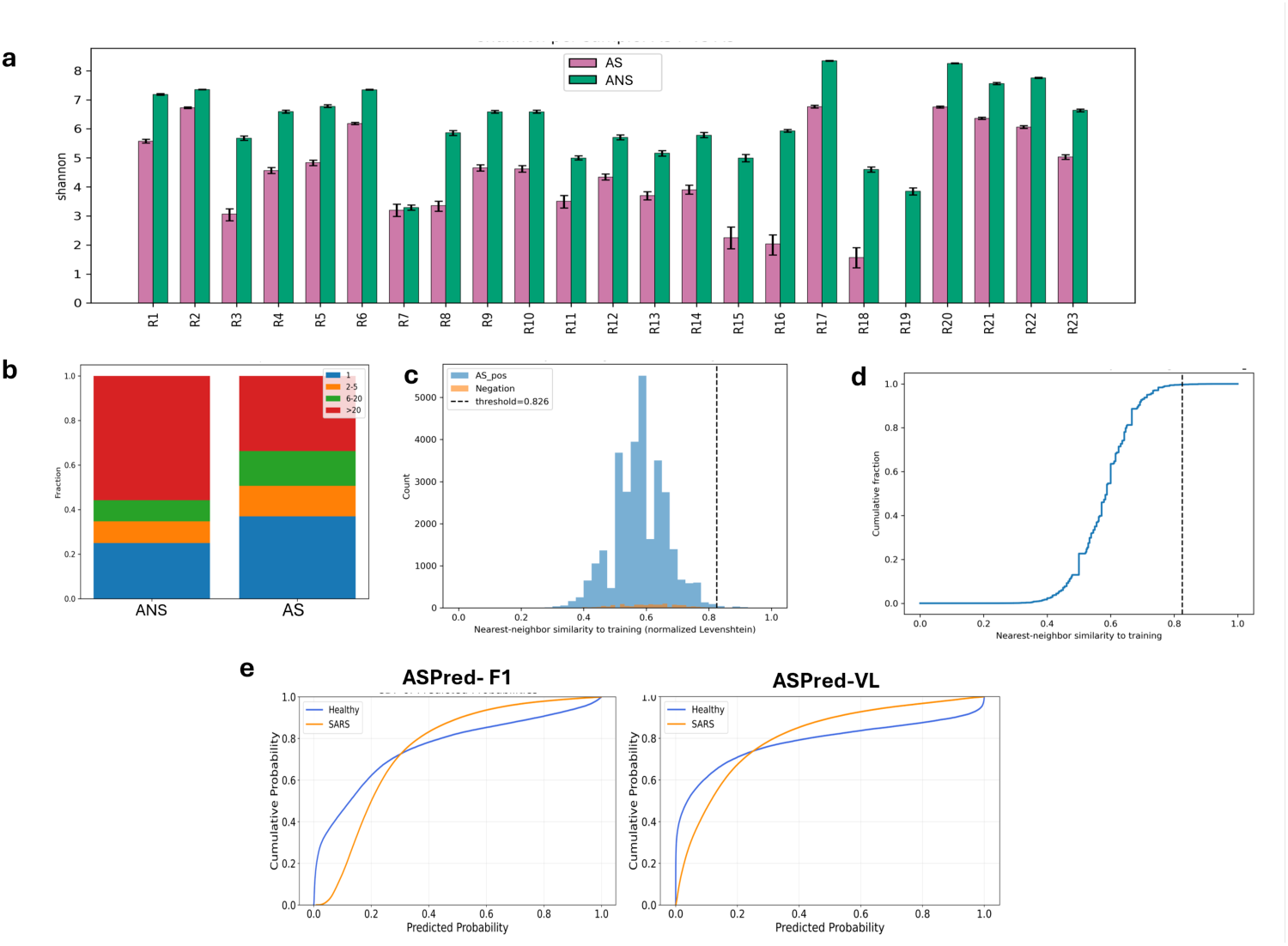
Diversity and training–data relatedness of ASPred^+^ BCRs in bulk repertoires. (**a**) Per-donor clonal diversity (Shannon entropy, *H*) for AS^+^ (magenta) and AS^−^ (green). Bars show mean with 95% bootstrap CIs (1000 resamples). (**b**) Novelty to training quantified by nearest-neighbor normalized Levenshtein similarity between each AS^+^ sequence and the training set (same-length CDRH3s); histogram shown with vertical dashed line indicating a negation-derived threshold (99th percentile from an unrelated influenza repertoire). (**c**) Empirical CDF of the same similarity measure for AS^+^; most points fall below the threshold, indicating low relatedness to training (i.e., novelty). (**d**) Clone-size composition (pooled across donors) for AS^+^ and AS^−^, showing the fraction of reads in bins 1, 2–5, 6–20, and *>* 20 members. (**e**) ROC curves quantifying separation of ASPred probability distributions between SARS-CoV-2–infected and healthy repertoires for the ASPred F1 and VL models.

### 2.9 ASPred predicted antigen-specific sequences do not resemble training data

To address whether AS^+^ migh resemble the training data, for each repertoire sequence we computed the normalized Levenshtein nearest-neighbor (NN) similarity to all sequences in the training dataset, and summarized the distributions with histograms and with empirical cumulative distribution functions (CDFs) (Figure 8c,d) [55]. The vast majority of AS^+^ sequences fell below a threshold of 0.8 NN similarity to the training set, indicating that ASPred’s predictions are not driven by near-exact sequence memorization. Finally, CDFs of ASPred SARS-CoV-2 class probabilities were right-shifted in the infected versus healthy repertoires, with the F1 model showing the largest separation (Figure 8f), consistent with antigen exposure.

## 3 Discussion

The identification of antigen-specific BCRs directly from complex immune repertoires is a fundamental challenge in immunology. In this study, we developed and validated ASPred, a transformer-based classifier capable of predicting antigen specificity directly from heavy chain sequences. By fine-tuning a large protein language model (ESM-2) on antigen-labeled antibody sequences and validating predictions computationally and experimentally, we demonstrate a scalable framework for identifying functional antibodies from whole immune repertoire sequencing data. While initially fine-tuned on SARS-CoV-2 RBD (nCoV-2019)-specific antibodies, the model maintained high predictive accuracy when applied to other viral antigens, including HIV gp120 and influenza hemagglutinin. These findings highlight ASPred’s potential as a broadly applicable framework for antigen-specific BCR classification, provided that well-labeled training datasets are available. Importantly, ASPred predictions were not only robust on held-out test sets but also were functionally validated through both *in silico* modeling and experimental binding assays.

From a set of ten antibody candidates selected randomly for experimental validation based on above-threshold ASPred scores, five demonstrated binding to the SARS-CoV-2 receptor-binding domain (RBD) by ELISA assays with strengths comparable to a commercial antibody used as a positive control. Among these candidates, Ab-157 was prioritized for detailed kinetic analysis and exhibited high-affinity binding to the RBD with a dissociation constant (*K*_D_) of 20.7 nM, a value sufficient for further development of a therapeutic antibody candidate.

The model’s ability to identify diverse binders that are phylogenetically distant suggests it is not merely memorizing sequence motifs, but is learning a deeper, underlying grammar of antibody function. We hypothesize that the model’s attention mechanisms may be capturing long-range electrostatic or hydrophobic interactions between framework residues and the CDR loops that are essential for presenting a binding-competent paratope. Furthermore, the model may be implicitly learning features of ‘developability,’ prioritizing sequences that are not only functional for binding but are also favored by cellular machinery for stable expression, thus providing a dual filter for both specificity and biophysical viability.

To benchmark ASPred’s performance, we compared it to BEAM, a widely used experimental method for determining repertoire-wide antigen-specific antibodies. While BEAM provides a useful framework for linking specificity to single-cell V(D)J sequences, it is limited by several practical and conceptual constraints. The requirement for highly purified antigens, which may lack native folding, introduces uncertainty, especially for antibodies where binding is frequently mediated by flexible CDR loops rather than rigid lock-and-key interactions. Moreover, BEAM is laborious and expensive, typically recovering only a limited number of V(D)J sequences from a small number of cells. By contrast, ASPred enables high-throughput analysis of thousands to hundreds of thousands of sequences *in silico*, allowing broader exploration of immune repertoire diversity and convergence patterns. When comparing ASPred’s predictions to BEAM-labeled sequences, we observed a representative overlap of sequences, which is consistent with the expectation that BEAM may fail to capture all true antigen-specific BCRs due to the limitations described above.

Conversely, ASPred may also miss certain binders if they are underrepresented or misrepresented in the training data, which are largely derived from curated antibody datasets such as OAS. These datasets tend to over-represent functional binders and lack comprehensive negative labels, *i.e.*, non-binding sequences that are inferred from the absence of signal, rather than from direct evidence of the lack of binding. Furthermore, the training data are not class-balanced; real repertoires contain far more non-binders than binders, a feature that is often underrepresented in fine-tuning datasets. Addressing this imbalance—through more realistic sampling, improved negative annotation, or semi-supervised learning—could improve classifier performance. Taken together, these considerations highlight both the complementary nature and inherent limitations of experimental and computational antigen mapping approaches.

Moving forward, integrating data from methods such as BEAM with broader immune repertoire sequencing may improve training of PLMs and further enhance the robustness of transformer-based predictors. Mining antigen-specific BCRs from human immune repertoires offers intrinsic safety and developability advantages, as these sequences have undergone natural immune tolerance and *in vivo* selection for proper folding and expression, thereby reducing the likelihood of immunogenic epitopes and aggregation-prone motifs [56].

When applying ASPred to bulk immune repertoire sequencing data from SARS-CoV-2-infected and healthy individuals, we observed clear shifts in the predicted probability distributions, despite the inherent heterogeneity and lack of baseline controls in such datasets. These shifts support the model’s capacity to extract informative signals from diverse B cell populations and suggest that, even in the absence of paired light chains or antigen pre-enrichment, the statistical signature of an antigen-specific immune response can be captured computationally.

We demonstrated by single-cell RNA sequencing that ASPred-predicted BCRs specific to the antigen were enriched among mature and lambda pre-B cell subtypes, suggesting that the classifier identifies biologically plausible candidates. Differential gene expression analysis further revealed transcriptional signatures associated with predicted binders, indicating that ASPred’s output corresponds to a functionally distinct B cell subset. Importantly, antigen-specific CDRH3 sequences did not form a single tight cluster in t-SNE or phylogenetic space, underscoring the model’s capacity to identify structurally diverse binders beyond simple clonal expansion patterns— an important feature for generalization. Nonetheless, these sequences had exhibited higher clonalities relative to nonspecific BCR sequences, as anticipated for antigen-stimulated clonal proliferation of B cells. By integrating repertoire-wide predictions with transcriptomic and structural analyses, we show that the model generalizes beyond clonal homology and captures underlying biophysical principles of antigen recognition. This work establishes a scalable and generalizable strategy for antibody discovery, with implications for vaccine design, immunotherapy, and synthetic antibody engineering.

In conclusion, ASPred provides a scalable and biologically grounded framework for the prediction of antigen-specific antibodies from bulk immune repertoires. Its integration with molecular simulation, transcriptomic context, and experimental validation offers a path forward for accelerating antibody discovery in both infectious disease and immunotherapy. Future work should focus on expanding antigen coverage, improving model interpretability using attention attribution and embedding analysis, and incorporating structure-aware sequence generation to design novel antibodies with desired biophysical and functional properties. Additionally, integrating paired light chain data, longitudinal repertoire tracking, and post-translational modification modeling (e.g., glycosylation) may improve prediction of context-specific binding and functional efficacy. Together, these efforts will enable more precise, data-driven interrogation of adaptive immunity and accelerate the discovery of next-generation biologics.

To facilitate broader use, we provide a public web interface for ASPred at https://ASPred.org/, enabling probability scoring of heavy-chain sequences without local computing.

## 4 Methods

### 4.1 Models and Data Sources

Two pretrained large language models were evaluated, namely ESM2-t6-8M and ESM2-t33-650M (facebook/esm2_t6_8M_UR50D, facebook/esm2_t33_650M_UR50D), for antigen specificity prediction. We attached a binary classification head to each backbone and tokenizing sequences with the corresponding ESM tokenizer. We used SARS-CoV-2 RBD–specific antibody heavy, light, or heavy-light sequences from CoV-AbDab and negatives sampled from PLAbDab (non–SARS-CoV-2 targets) ([24], [57]). Model training was performed using 3-fold cross-validation, and the best configurations were retrained on the full datasets before evaluation on held-out test sets. Per-fold performance summaries are shown in Extended data, and full configuration details Supplementary Table 1.

### 4.2 Identification of antigen-specific BCRs using large language models

#### Initial fine-tuning

We performed an end-to-end fine-tune of the 6-layer ESM-2 8M model (facebook/esm2_t6_8M_UR50D) using a single-label classification head (EsmForSequenceClassification). Inputs were tokenized with the ESM tokenizer (EsmTokenizer; vocabulary size 33) and padded/truncated to 1,024 tokens. Model hyperparameters followed the saved configuration (config.json): hidden size 320, intermediate size 1,280, 20 attention heads, GELU activations, layer norm *E* = 10*^−^*^5^, hidden dropout = 0.1, attention dropout = 0.0, rotary position embeddings (max positional capacity = 1,026), and output_hidden_states=true; training used FP32 under transformers v4.32.1. Trained model is provided as Extended Data File 1

#### Parameter-efficient fine-tuning (PEFT/LoRA)

The fine-tuning used LoRA adapters inserted into each block’s self-attention projections (W*_q_,* W*_k_,* W*_v_*), with all other backbone weights frozen and the classification head trainable. We ran Optuna hyperparameter optimization under stratified 3-fold cross-validation in two complementary modes: first, an F1-maximization study with search ranges lora_r *∈* {4, 8}, lora_alpha *∈* [16, 64], lora_dropout *∈* [0.05, 0.10], learning rate [5 *×* 10*^−^*^7^, 2 *×* 10*^−^*^5^] (log-uniform), batch size {4, 8}, epochs [4, 8], and warmup ratio [0.05, 0.20], trained via transformers.Trainer with early stopping (patience=1), fp16 when CUDA was available, gradient accumulation (2 steps), pinned dataloaders, and selection by mean F1 across folds; second, an explicit validation-loss minimization study that optimized eval_loss by setting metric_for_best_model=eval_loss and greater_is_better=False, widened certain ranges (lora_r *∈* {4, 8, 16}, lora_dropout *∈* [0.05, 0.30], learning rate [10*^−^*^6^, 5 *×* 10*^−^*^5^], epochs [2, 6], warmup ratio [0.05, 0.20], batch size {4, 8}), and returned the negative average fold loss to keep a maximize objective. Both studies logged per-fold accuracy, precision, recall, F1, ROC–AUC, confusion matrices, ROC curves, and artifacts to Weights & Biases with timestamped project names, and after HPO we retrained the selected backbone on the full dataset with best hyperparameters, exported PEFT-augmented checkpoints, and saved Optuna studies (.pkl) plus optimization-history and parameter-importance plots under a versioned directory structure; evaluation reported F1 as primary along with accuracy, precision, recall, ROC–AUC, and confusion matrices computed with scikit-learn, deriving ROC from softmax positive-class probabilities and writing per-fold predictions (label, pred, prob) to CSV. Jobs were executed on an HPC cluster under Slurm using L40S GPUs (typically 2*×* L40S, 120 GB RAM, 20 CPU cores) with environment modules gcc/13.3.0, mpich/4.2.2, and a Conda environment containing pytorch, transformers, peft, optuna, and wandb.

### 4.3 BEAM - ASPred overlap

#### Dataset

We first tested ASPred’s predictions on V(D)J heavy chain sequences with BEAM specificity scores against the RBD antigen as a probe against B-cells isolated from blood from SARS-CoV-2 infected human subjects (whole PBMCs, Single Cell BEAM Ab Immune Profiling Dataset by Cell Ranger v3.1.0, proprietary data provided by 10x Genomics). We determined the overlap of positive classified sequences by ASPred and BEAM (Extended Data File 2). The same BEAM dataset contained a coupled-barcoded scRNA-seq gene expression data between antigen-specific (AS) from non-specific (AS) B cells, which allowed us to determined differential gene expression levels between them using the Seurat (v5.0.1) package, and the Wilcoxon rank-sum test. Cells were annotated by SingleR.

### 4.4 Ethics statement

All animal procedures were approved by the University of California, Riverside Institutional Animal Care and Use Committee (IACUC) and conducted in accordance with institutional and federal guidelines (protocol #AUP20210029).

### 4.5 Immunization and sample collection

Mice were housed at the University of California Riverside vivarium. BALB/c female mice at 6 weeks of age were given subcutaneous injections of 100 *µ*L per day in the back of the neck on days 0 and 14 of antigens emulsified in aluminum hydroxide 2 % (RBD protein SARS-CoV-2 2019-nCoV). Blood samples were collected post-immunization and on days 14 and 28. On day 28 mice were deeply anesthetized with isoflurane, and blood was drawn by cardiac puncture. Mice were immediately euthanized by cervical dislocation according to IACUC guidelines. Peripheral Blood Mononuclear Cells (PBMCs) were isolated using the direct human PBMC isolation kit (StemCell Technologies) and cryopreserved at -80*^◦^*C for further work.

### 4.6 GEM Generation and construction of gene expression next-generation sequencing libraries

PBMCs were used without antigen labeling and sorting. Cryopreserved peripheral blood mononuclear cells (PBMCs) from immunized mice were thawed rapidly at 37°C, transferred to pre-warmed complete growth medium, and centrifuged at 300 × g for 5 minutes to pellet cells. The cell pellet was resuspended in PBS supplemented with 0.04% BSA. Cell concentration and viability were determined using a manual hemacytometer with trypan blue exclusion, ensuring viability exceeded 85%). Cells were then adjusted to a final concentration of 700–1,200 cells/µL. Single-cell suspensions were mixed with nuclease-free water and 5*^1^* single-cell RNA master mixture, then loaded into a Chromium chip with barcoded gel beads and partitioning oil. The chip was placed in the Chromium controller to generate gel beads in emulsion (GEMs). cDNA was obtained from 100 *µ* l GEMs/sample by reverse-transcription reactions: 53 *^◦^* C for 45 min, 85 *^◦^* C for 5 min, then maintained at 4 *^◦^*C. cDNA products were purified and cleaned using Dynabeads (Derbois et al., 2023). cDNA was amplified by PCR: 98*^◦^*C for 45s; 98 *^◦^*C for 20 s, 63 *^◦^*C for 30 s, 72 *^◦^*C for 1 min and amplified for 16 cycles; then, 72 *^◦^*C for 1 min. Amplified PCR products were purified using SPRIselect reagent kit (B23317, Beckman Coulter). The concentration of the cDNA library was determined by Qubit dsDNA HS Assay Kit (Invitrogen) and Bioanalyzer (Agilent, 2100) (Wang et al, 2023). Single Cell RNA-Seq V(D)J and 5’ gene expression library was performed using the Chromium Next GEM Single Cell 5’ Reagent Kits v2 (Dual Index)(CG000331, Rev E, 10X Genomics) and Dual Index kit TT set A (PN1000215, 10X Genomics) according to the manufacturer’s instructions. For unlabeled and unsorted samples, the target was estimated at 5000 cells.

### 4.7 Identification of antigen-specific BCRs using interclone

Clustering was performed using the source code of InterClone, which employed MMSeqs2 for clustering. The dataset being clustered was constructed with 11,917 known SARS-CoV-2 specific antibody heavy chain sequences from CovAbDab [58] and 310 heavy chain sequences from the single cell sequencing data. To identify antigen-specific B cell receptors (BCRs) from the total repertoire of peripheral blood mononuclear cells (PBMC), we employed three distinct approaches. The first approach utilized the clustering tool *InterClone* [59], which introduces a novel method to cluster antibodies sharing antigenic targets based on their complementarity-determining region (CDR) sequences, with *MMSeq2* facilitating effective sequence clustering through homology alignment and gap management. We constructed a dataset comprising 11,917 known SARS-CoV-2-specific antibody heavy chain sequences sourced from *CovAbDab* [58] and an additional 310 heavy chain sequences obtained from our own experimental single-cell testing dataset. Using a clustering threshold of 70% for the CDR similarity index (SID) and requiring 90% coverage, we identified 96 candidate sequences from our single-cell testing dataset, which clustered with known SARS-CoV-2 antibody sequences.

### 4.8 Cloning and recombinant VHH expression

DNA molecules of VHH sequences were synthesized by Integrated DNA Technologies (IDT). Each 20 ng fragment was PCR-amplified using a New England Biolabs (NEB) kit. The pET-22b(+) vector (100 ng) was linearized with NotI and NcoI, heat-inactivated at 80*^◦^*C for 20 min, and assembled with 50 ng of PCR-amplified fragments by Gibson assembly using the NEBuilder^®^ HiFi DNA Assembly Master Mix according to the manufacturer’s instructions (50*^◦^*C for 60 min). *E. coli* DH5*α*; transformants were screened by colony PCR and confirmed by Sanger sequencing. *E. coli* BL21 (DE3) competent cells were transformed with the plasmid constructs. Single colonies were cultured at 37*^◦^*C to an OD_600_ of 0.6, induced with 1 mM isopropyl *β*-D-1-thiogalactopyranoside (IPTG) at 37*^◦^*C for 4 h, and then incubated with agitation at 22*^◦^*C overnight. Cultures were centrifuged, and cell pellets were resuspended in phosphate-buffered saline (PBS; 137 mM NaCl, 2.7 mM KCl, 10 mM Na_2_HPO_4_, 1.8 mM KH_2_PO_4_, pH 7.4). Proteins were extracted by freeze–thawing twice. Samples were dialyzed overnight in 1 L of PBS at 4*^◦^*C. Protein expression was confirmed by dot blot using an HRP-conjugated anti-His antibody (Ay63, Cat. HIS-PLM535). As a positive control, a His-tagged S1S2 protein was used; negative controls included *E. coli* BL21 cells cultured without plasmid and Q*β* protein. Western blot analysis was performed by probing with the anti-His antibody; extracts of BL21 cells without plasmid served as the negative control. All constructs were expressed in *E. coli*, and a freeze–thaw protocol previously shown to efficiently release nanobodies into the culture supernatant [60] was used to obtain nanobody-rich supernatants for screening.

### 4.9 Analysis of antigen immunogenicity in mice by Enzyme Linked-Immunosorbent Assay (ELISA)

To evaluate antigen binding, we performed an initial ELISA screen against the SARS-CoV-2 receptor-binding domain (RBD) using these unpurified supernatants. This assay aimed to detect any binding activity regardless of antibody concentration. ELISA was carried out in 96 half-area well plates from (Greiner Bio-One), plates were coated with the full-length spike protein from SARS-CoV-2 nCov-2019 (0.4 *µ* g/well) using sodium carbonate coating buffer (0.05M, pH 9.6) and allowed to incubate overnight at 4 ° C (Pan et al. 2021). Plates were washed (2x) with PBST (PBS with 0.05% Tween-20) and once with PBS. Plates were blocked with a 5% non-fat dry milk solution for 1 hour. Samples were added (dilutions: 1:5), and each dilution was incubated with immobilized antigens for 1 hour at room temperature with continuous shaking (Ye et al., 2021). The plates were washed 5 min each 2x with PBST and 1x with PBS. The bound VHHs were then detected with anti histidine antibody diluted at 1:2,500 in PBS by incubating at RT for 1 hr, and the plates were washed 2x in PBST, rinsed 1x with PBS. The enzymatic reaction was initiated by adding the OPD Peroxidase substrate (Sigma Alrich, P9187-50SET) to the wells and allowing the reaction to proceed for 5 minutes and the developed color was measured using an ELISA microplate reader (NanoQuant Infinite M200, Tecan) at 450 nm after 5 and 30 minutes (Alsoussi et al., 2020).

### 4.10 Localized surface plasmon resonance experimental binding validation

LSPR measurements were performed on a Nicoya OpenSPR-XT instrument using a High-Capacity Carboxyl sensor (Nicoya) and a PBS-T pH 7.4 running buffer (1*×* PBS, 0.05% Tween-20). Binding experiments were performed at room temperature. Analyte VHH samples were diluted to the indicated concentrations in PBS-T. RBD was used as the ligand immobilized to the carboxyl sensor and Ab-157 was used as the analyte in solution. Carboxyl sensor surfaces were activated with 10 mM 1-ethyl-3-(3-dimethylaminopropyl) carbodiimide (EDC) and N-hydroxysuccinimide (NHS) in ultrapure water (MilliQ, resistivity 18.2 MΩ·cm). RBD was prepared at 50 *µ*g/mL in sodium acetate pH 5.0 and coupled to the surface to *∼*1800 RU at a flow rate of 10 *µ*L/min. RBD immobilization gave a stable immobilization response with minimal surface degradation over time. Unreacted NHS esters were blocked with 1 M ethanolamine prior to analyte injection. Binding and unbinding experiments were run at a flow rate of 40 *µ*L/min. Between each measurement cycle, the sensor surface was regenerated with two 10-second injections of 10 mM Glycine-HCl pH 2.0 to remove bound Ab-157 from the RBD surface. A 1:1 binding model with simultaneous curve fitting was applied. The kinetic results were analyzed with the manufacturer suggested analysis in TraceDrawer 1.9.1. The analysis included subtraction of the reference channel from the immobilized active channel of RBD.

### 4.11 Prediction of antibody structure using Igfold and Docking of antibody candidates with RBD

Variable heavy chain protein sequences obtained by ASPred were folded using IgFold [44] and the structures were refined with PyRosetta [61] Antibody sequences were renumbered according to the Chothia scheme [62]. The RBD protein sequence was folded using AlphaFold 2 [63] and used as ligand, while antibody structures served as receptors in the ClusPro 2.0 docking web server [64, 65] with the antibody mode and non-CDR masking options enabled

### 4.12 Binding affinities for the complexation of antibody candidates with RBD

The Fast cOarse-grained pRotein-proTein modEl (FORTE) [48, 49] is a coarse-grained biophysical model specifically designed to simulate protein-protein interactions, allowing for the dynamic adjustment of amino-acid charges on titratable groups based on the surrounding environment at a specified pH (input as a parameter). The core of this model is the FPTS [51] combined with the ability to translate and rotate macro-molecules using the Metropolis MC method [66] All calculations with FORTE were performed at pH 7.4, 150 mM NaCl, and 298K. For a more accurate incorporation of hydrophobic interactions, we used a Lennard-Jones potential energy with *ε_LJ_* equivalent to 0.1736 kJ/mol. All other methodological details and parameters are provided in reference [48]. This model enables molecular simulations that are computationally faster than more complex models, providing relative binding affinities at a reduced computational cost. This allows for the comparison across various molecular systems and/or physicochemical conditions. The heavy chains of the top candidates predicted by ASPred were folded using IgFold and then used as input for the FORTE simulations. To ensure comparability with previous studies [49, 50, 67] the wildtype RBD structure was constructed via the SWISS-MODEL workspace, using the NCBI reference sequence NC_045512 (accession YP_009724390.1) as a template. This approach facilitated a direct comparison of the binding affinities between ASPred-predicted antibodies and previously characterized ones [49, 50] The free energies of interaction (or binding affinities) were calculated as a function of the macromolecules’ separation distances by analyzing their center-to-center pair radial distribution functions. These values were sampled in histogram form during the MC production phase, providing detailed distributions of the probability of finding the two molecules at different separation distances. To compare binding affinities across systems, we adopted the free energy minima (*βA*) observed in these simulations [49, 50], which represent the most stable interaction points. This approach is a simple and consistent basis for evaluating relative affinities between the top candidates.

Following the equilibration phase, each system underwent at least 3 × 10^9^ MC steps for production-phase sampling. To account for variability, three independent replicates were conducted for each system, allowing for the estimation of statistical errors for a proper comparison between the simulated systems.

### 4.13 Electrostatic stability of antibody candidates

The electrostatic stabilities of the antibody heavy chains were calculated directly from the averaged net charges obtained in the FPTS simulations, using the IgFold-generated conformations [68, 69]. Coulombic contributions of individual titratable groups for each protein structure were measured.

### 4.14 Molecular dynamics simulations of antibody candidates with RBD

Molecular dynamics simulations were performed using GROMACS 2024.3 with the AMBER99SB-ILDN protein force field and the TIP3P water model. The antibody– RBD complex was placed in a cubic box with at least 0.2 nm clearance from the edges, solvated with water, and neutralized with 0.15 M NaCl. Energy minimization was followed by equilibration at 300 K and 1 bar for 100 ps each in the NVT and NPT ensembles, respectively. Production simulations were run for 300 ns with a 2 fs integration step. Nonbonded interactions were calculated with a cut-off of 1.0 nm for both Coulomb and van der Waals interactions, with long-range electrostatics treated by the particle-mesh Ewald method. Temperature was maintained using the V-rescale thermostat (*τ* = 0.1 ps) applied separately to protein and solvent, and pressure was controlled using the Parrinello–Rahman barostat (*τ* = 2.0 ps, compressibility 4.5 *×* 10*^−^*^5^ bar*^−^*^1^). Bond lengths involving hydrogens were constrained with the LINCS algorithm. Trajectories were analyzed after removal of periodic boundary conditions.

### 4.15 Diversity of BCRs and sequence similarity to training data

To assess whether predicted antigen-specific BCRs (ASPred^+^) represented previously unobserved solutions or were closely related to sequences in the training set, we quantified their sequence-level novelty using a nearest-neighbor (NN) similarity framework based on normalized Levenshtein distances [70]. Briefly, CDRH3 amino acid sequences from the ASPred^+^ subset were compared to all antigen-specific heavy-chain sequences used for model training. For each query sequence *q_i_*, the highest normalized Levenshtein similarity to any training sequence *t_j_*was computed as:

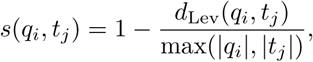

where *d*_Lev_(*q_i_, t_j_*) denotes the standard Levenshtein edit distance between the two sequences, and *s*(*q_i_, t_j_*) *∈* [0, 1] corresponds to identical sequences when *s* = 1. The nearest-neighbor similarity for a query was defined as *S_i_*= max*_j_ s*(*q_i_, t_j_*). Computations were performed using the RapidFuzz implementation for efficiency. When specified, comparisons were restricted to same-length CDR3s to approximate Hamming similarity. A novelty threshold *T* was defined empirically using a “negation” reference set comprising antigen-unrelated or ASPred^−^ BCRs. The nearest-neighbor similarities of these negation sequences to the training set were computed, and *T* was chosen as the (1 *− δ*)-quantile of their similarity distribution, corresponding to a false-positive tolerance *δ* = 0.01:

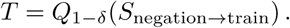

ASPred^+^ sequences with best NN similarity *S_i_ <T* were considered *novel*. Per-donor novelty frequencies were summarized as the fraction of ASPred^+^ BCRs classified as novel, with 95% Wilson confidence intervals [71]. Histogram and cumulative distribution plots were used to visualize similarity distributions and the defined threshold.

### 4.16 Phylogenetic analysis of antigen specific sequences

We started with three datasets, and found the positive intersection of ASPred and BEAM with probability higher than 0.5 from them. This intersection yielded 127 sequences. ANS sequences from healthy human repertoire were considered negative or antigen nonspecific. We took the 127 positive sequences and generated 200 separate fasta files that include the same 127 positive sequences and 127 randomly selected sequences from the two negative datasets. These 200 mulifastas were then aligned using mafft using the auto settings. Then we ran RAxML-NG (MPI) to run a ML tree search using the specified protein model LG+G4+F (LG exchangeability matrix, 4-category discrete Gamma rate heterogeneity, empirical AA frequencies), and used standard nonparametric bootstrapping with 100 replicates. We took the best trees and bootstraps from each run and then computed Fitch parsimony for AS/ASN on the tree and a permutation null (n=999). We also recorded the sizes of AS-only clades (all descendants AS) of size 3. Finally, we measured local AS enrichment by taking the mean fraction of AS among k=3 nearest neighbors

## Acknowledgements

This work was funded in part by the National Institute of Allergy and Infectious Diseases (NIAID) of the US National Institutes of Health (NIH) under the award 3R01AI169543 to S.L., M.S., J.B.H., and A.R., and also was supported in part by the Fundação de Amparo a’ Pesquisa do Estado de São Paulo (Fapesp) [Grant 2020/07158-2 to F.L.B.d.S.] and the Conselho Nacional de Desenvolvimento Científico e Tecnológico (CNPq) [Grant 305393/2020-0 to F.L.B.d.S.]. A.R. acknowledges access to the Center for Advanced Research Computing infrastructure LAGUNA HPC cluster of the University of Southern California through the SoCal Research Computing Alliance; F.L.B.d.S. acknowledges the Swedish National Infrastructure for Computing (SNIC) at NSC and PDC for providing resources used to run the constant-pH simulations; S.L. acknowledges the High Performance Computing Center at University of California Riverside for the training of ASPred.

## Declarations

### Code availability

All scripts used for parameter optimization of ESM-2 models with Optuna and Weights and Biases are available via GitHub at github.com/karenpaco/ASPred_raylab. The repository includes multiple optimization strategies (e.g., F1 score with early stopping, and validation loss). To replicate the results, add your data to the datasets/directory and adapt the objective function as needed.

### Author contribution

K.P. conceived the study, designed analyses, developed methodology, coordinated the project, guided experimental work, performed computational analyses, prepared figures and wrote all drafts of the manuscript. A.R. provided overall supervision and guidance, contributed to study conception, contributed analysis, reviewed and edited the manuscript, and secured resources. Z.Z. developed the first version of the model; M.P.-M., Z.Z., and J.A.L. implemented model training and performed molecular-dynamics simulations. F.B.A. implemented and analyzed the ABLang component. K.P., P.O., I.C.R., and J.H immunized mice and perform single cell analysis. K.P., C.D. performed molecular docking. T.Y. performed sequence clustering and phylogenetic analyses. S.Z., R.M.A., P.O., I.C.R. and E.C. carried out wet-lab experiments, purified recombinant proteins and carried out in vitro analysis. K.P., R.M.M. performed LSPR measurements. F.L.B.S. conducted coarse grained simulations and contributed to the conception of the work and to manuscript revisions. S.L., J.B.H., M.H.S., J.F., and I.T. contributed to study conception and provided critical feedback. V.G. and S.B. developed the ASPred web server. K.L.R. and M.S.J. supported instrumentation and execution of 10x sequencing and SPR experiments respectively and advised on experimental design. K.P. and A.R. verified the overall integrity of the work. All authors discussed the results and approved the final version of the manuscript.

#### Equal contribution

M.P.-M., Z.Z. and S.Z. contributed equally to this work.

