## Supplemental Information for "Fine-tuned protein language model identifies antigen-specific B cell receptors from immune repertoires"

2Pomona College, Claremont, CA, USA.

5Universidade de São Paulo, Brazil.

6Department of Chemical and Biomolecular Engineering, NC State University, Raleigh, NC, USA.

7Toyoko Labs, Toyoko LLC, Emeryville, CA,USA.

8Division of Biology and Biological Engineering, California Institute of Technology, Pasadena, CA,USA.

**SUPPLEMENTARY RESULTS 1: Model Development.** Full-parameter fine-tuning of the 6-layer ESM-2 8M backbone (`esm2_t6_8M_UR50D`) with a single-label sequence-classification head on SARS-CoV-2 RBD heavy-chain sequences (Model 1) yielded a classification accuracy exceeding 80% (Figure S1), versus ∼60% for the pretrained baseline. This direct fine-tuning established feasibility, and motivated a broader evaluation across architectures and input modalities. To enable efficient and systematic fine-tuning, we applied low-rank adaptation (LoRA) to the ESM-2 architecture. Supplementary Table 1 summarizes the model configurations, and descriptions of training datasets are provided in Extended Data, and their partitions are diagrammed in Figure 1 and in Methods.

To evaluate model performance across different sequence inputs and antigen targets, we leveraged three sequence types: heavy-only, light-only, and stacked (heavy + light). We compared two model sizes—8M and 650M parameters—across three different antigen types: the SARS-CoV-2 Spike protein's receptor binding domain (SARS-CoV-2 RBD (2019-nCoV)), HIV gp120, and influenza hemagglutinin (HA). Hyperparameters were optimized using Optuna [72] with two separate objectives: maximizing F1-score and minimizing validation loss (Supplementary Figure S4, and Supplementary Table 1). We refer to the resulting best models as Antigen Specificity Predictor F1-optimized (ASPred F1) and validation-loss–optimized (ASPred VL).

The fine-tuned ESM-2 model surpassed 80% accuracy on the test dataset (Figure S2). By contrast, a top-of-the-line protein language model pre-trained on a large corpus of antibody sequences *AbLang* [31], used to generate sequence embeddings and supplied to a logistic regression classifier, achieved an accuracy of 55.2% (±0.0091) on the same dataset (Extended Data Fig. 1). Therefore, while antibody-specific models such as AbLang may perform well in restoration tasks [31], predicting a complex functional phenotype, such as antigen-specificity from a whole immune repertoire, benefits more from the representational capacity of a model fine-tuned on a task-specific dataset.

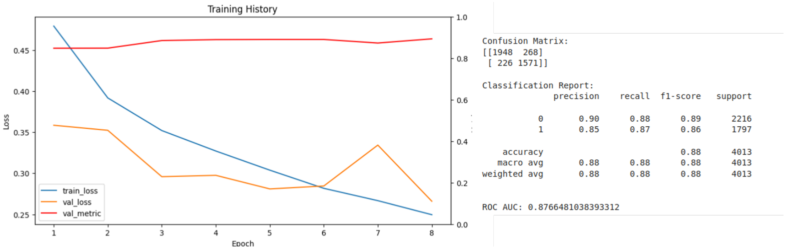


**Fig. S1**: ESM-2 fine tuned model, trained on SARS CoV-2 specific heavy chain sequences. Model 1 test performance. Confusion matrix and classification report on the held-out test set: AUROC = 0.877, accuracy = 0.88; TN = 1948, FP = 268, FN = 226, TP = 1571

**BEAM-ASPred Ovelap Estimation.** Across BEAM-Ab assays, ASPred probabilities tracked BEAM scores: median BEAM increased monotonically by ASPred deciles, and top-$q$ selections (1%, 5%, 10%) showed enriched overlaps with BEAM top-$q$ (elevated Jaccard and mutual coverage). Using BEAM $\geq$80 as a “high” threshold, ranking by ASPred markedly improved early hit rates over a random baseline; results were consistent when sweeping BEAM cutoffs (70, 80, 90) and when comparing both classifiers (ASPred-F1 and ASPred-VL).

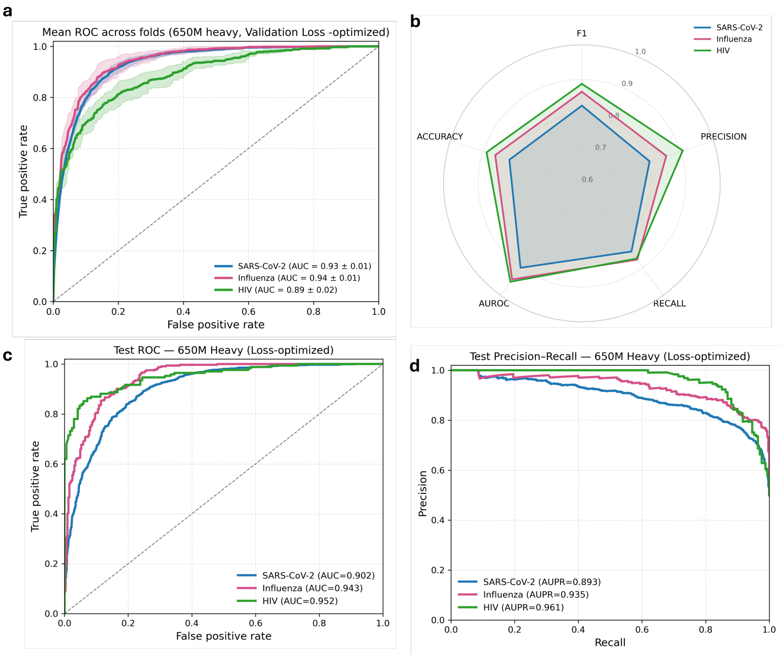


**Fig. S2**: ASPred VL Receiver Operating Characteristic (ROC) curve, )Precision Recall (PR) and Radar plot summarizing performance metrics for the final model ASPred-VL, evaluated on an independent held-out test set.

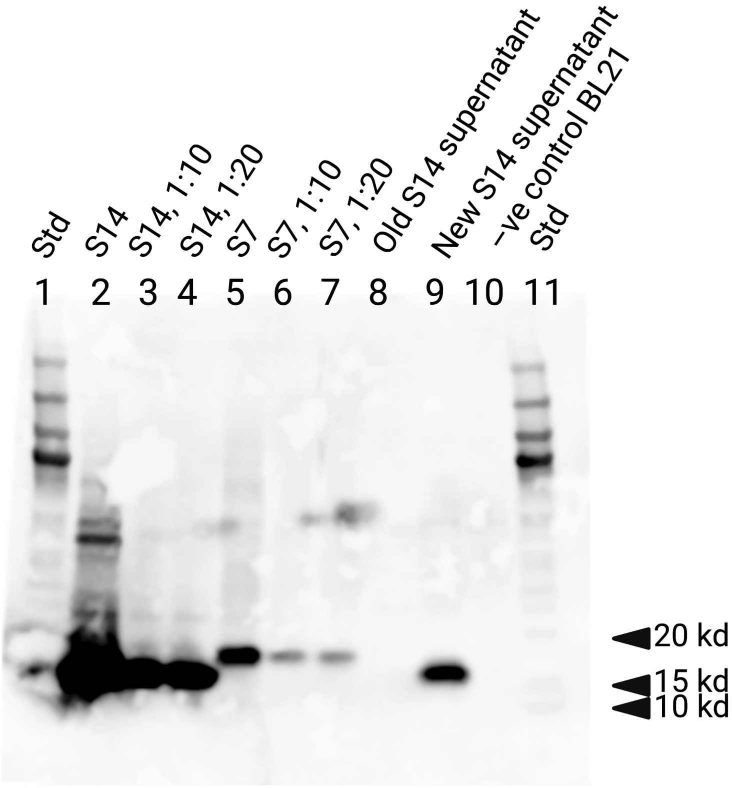


**Fig. S3**: Western Blot purified VHHs Ab-157 (S7) and positive control VHH 7OAO (S14)

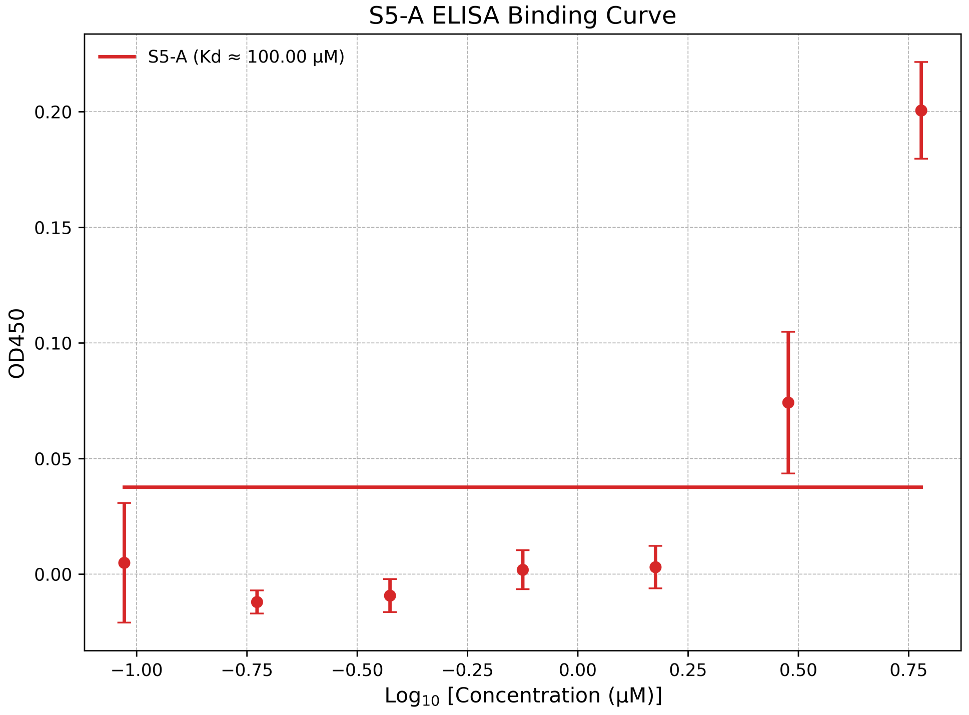


**Fig. S4**: Purified Ab-200 (S5) shows a concentration-dependent increase in absorbance (OD450) when tested against immobilized SARS-CoV-2 (2019) RBD, indicating binding above background at higher concentrations. Points are mean ± s.e.m. (n=3); the solid line is the fit; the horizontal line marks background.

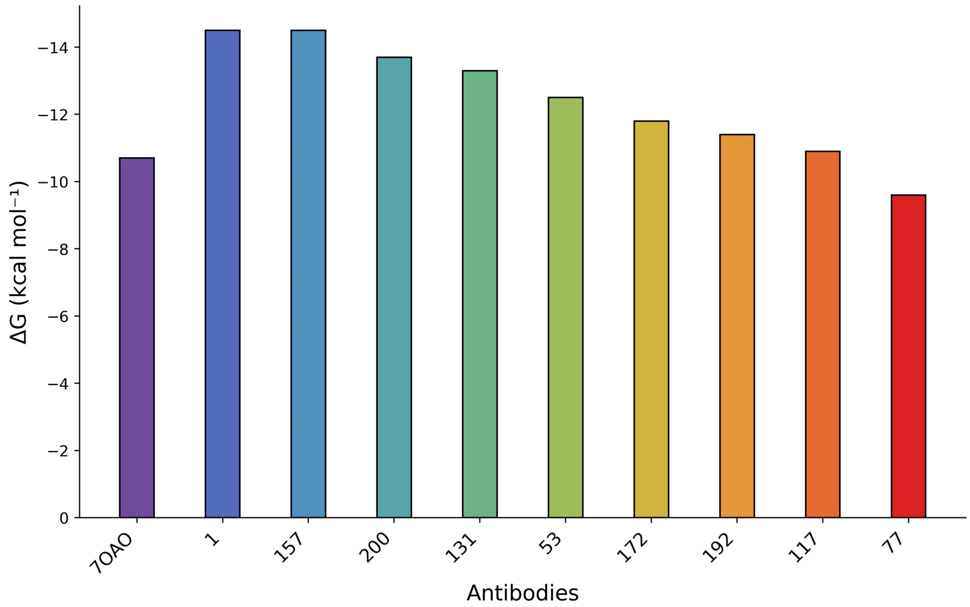


**Fig. S5**: Docking ClusPro and binding affinity prediction by PRODIGY.

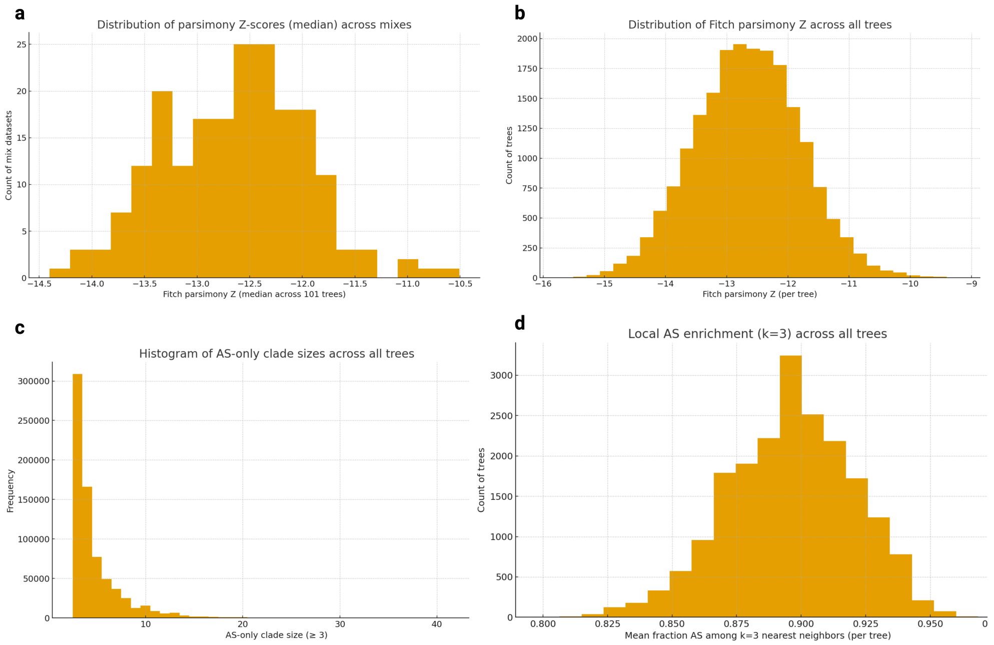


**Fig. S6**

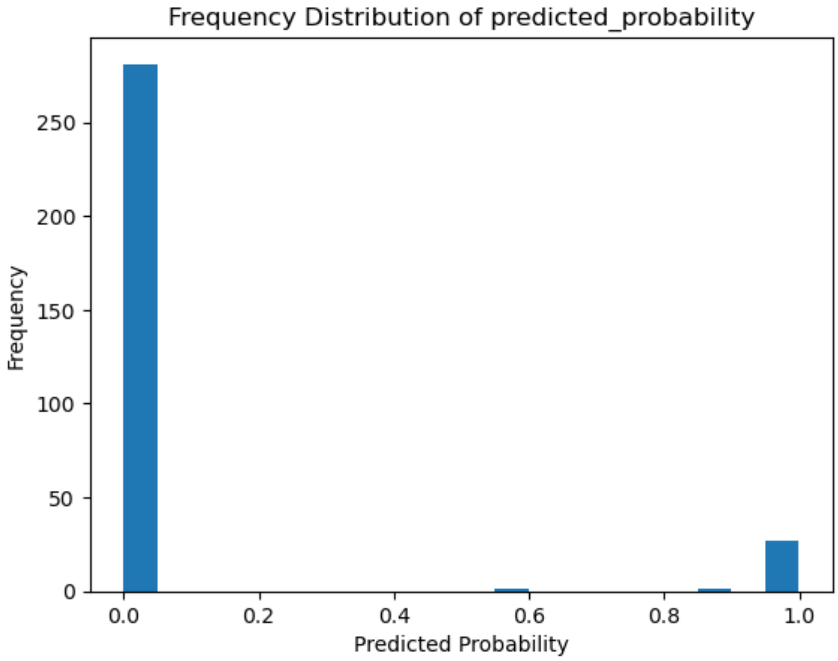


**Fig. S7**: ASPred Model 1 frequency distribution of probability scores for V(D)J sequences from RBD-immunized PBMC immune repertoire.

**Supplementary Table 1**: **Summary of language models used in this study.** Each row corresponds to an ESM-2 model variant fine-tuned for antibody specificity prediction. *Encoder Params* indicates the total number of trainable parameters in the encoder; *LoRA Params* represents the number of trainable parameters introduced through Low-Rank Adaptation (LoRA); *Emb Dim* denotes the embedding dimensionality of token representations; and *HuggingFace Checkpoint* refers to the original pretrained model identifier used for fine-tuning.

| **Model** | **Architecture** | **Encoder Params** | **LoRA Params** | **Layers** | **Embedding Dim** | **HuggingFace Checkpoint** |
|---|---|---|---|---|---|---|
| ESM2-8M | Encoder-only Transformer | 8 M | 163 K | 6 | 320 | `esm2_t6_8M_UR50D` |
| ESM2-650M | Encoder-only Transformer | 650 M | 3.5 M | 33 | 1280 | `esm2_t33_650M_UR50D` |
